# Specific configurations of electrical synapses filter sensory information to drive choices in behavior

**DOI:** 10.1101/2023.08.01.551556

**Authors:** Agustin Almoril-Porras, Ana C. Calvo, Longgang Niu, Jonathan Beagan, Josh D. Hawk, Ahmad Aljobeh, Elias M. Wisdom, Ivy Ren, Malcom Díaz-García, Zhao-Wen Wang, Daniel A. Colón-Ramos

## Abstract

Synaptic configurations in precisely wired circuits underpin how sensory information is processed by the nervous system, and the emerging animal behavior. This is best understood for chemical synapses, but far less is known about how electrical synaptic configurations modulate, *in vivo* and in specific neurons, sensory information processing and context-specific behaviors. We discovered that INX-1, a gap junction protein that forms electrical synapses, is required to deploy context-specific behavioral strategies during *C. elegans* thermotaxis behavior. INX-1 couples two bilaterally symmetric interneurons, and this configuration is required for the integration of sensory information during migration of animals across temperature gradients. In *inx-1* mutants, uncoupled interneurons display increased excitability and responses to subthreshold temperature stimuli, resulting in abnormally longer run durations and context-irrelevant tracking of isotherms. Our study uncovers a conserved configuration of electrical synapses that, by increasing neuronal capacitance, enables differential processing of sensory information and the deployment of context-specific behavioral strategies.

**One-Sentence Summary:** Coupling of interneurons by electrical synapses reduces membrane resistance and filters sensory inputs to guide sensory-dependent behavioral choices.

## Main Text

Behavioral outputs rely on sensory information. Sensory information can be differentially processed based on the configurations of synapses in the circuit, enabling *similar* sensory stimuli to elicit *different* behavioral strategies in context-dependent manners (*1–15*). This action selection (*16–18*) enables animals to avoid deploying incompatible locomotory strategies in response to similar sensory stimuli at behavioral choice points. While the importance of action selection in behavioral choice strategies is well-recognized (*16–18*), the synaptic configurations that support action selection are not well understood.

Dissecting action selection mechanisms at a circuit level requires: 1) deriving predictable choice points for a given behavioral paradigm, 2) knowing the circuit substrates underlying the behavioral choice points and 3) understanding sensory input processing and locomotory strategy selection at the behavioral choice points. *C. elegans* thermotaxis behavior (*19*) provides a tractable model to interrogate the circuitry and synaptic bases of action selection*. C. elegans* does not have an innate preferred temperature, and instead learns to prefer the temperature at which it was cultivated in the presence of food (*19*). When in a temperature gradient, animals perform two behavioral strategies to reach and stay within their learned preferred temperature: migration across the gradient to arrive at the previously experienced temperature range (gradient migration), and tracking of isotherms upon encountering their preferred temperature (isothermal tracking) (*19*). Gradient migration and isothermal tracking are two behaviors that cannot be performed simultaneously. Because the action selection switch between gradient migration and isothermal tracking occurs within the temperature window at which the animal was cultivated, thermotaxis behavior provides an assay in which the behavioral choice point is both predictable and quantifiable. Importantly, the specific neurons that underlie thermotaxis behavior have been identified (*11, 20–28*). Laser-ablation studies of neurons in this circuit produces defects in both isothermal tracking and gradient migration (*11, 20–24, 29, 30*), indicating shared circuitry between the two strategies. How synaptic configurations in this circuit influences processing of thermosensory information to deploy context-specific behavioral strategies is not known.

To uncover circuits that underpin action selection mechanisms, we performed behavioral genetic experiments in *C. elegans*. We first adapted a thermotaxis assay to enrich for the quantification of isothermal tracking and gradient migration in a population of isogenic animals (Fig. 1 and Supp. Fig. 1). Animals were placed in separate regions of a temperature gradient with regards to their preferred temperature goal (20°C), and the locomotory strategies were recorded, segmented and quantified while they performed gradient migration and isothermal tracking (Fig. 1). Consistent with previous reports (*31*), under these conditions wild-type *C. elegans* spent about ∼12% of their total time on the assaying arena performing isothermal tracking when within ±2°C of their preferred temperature, and with an average duration of 65 seconds per isothermal-oriented run (Fig. 1B, E). Distribution of the durations of isothermal track events followed an exponential decay with a time constant of 49.61 seconds and a half-life of 34.39 seconds (Fig. 1G and (*31*)).

**Figure 1.**
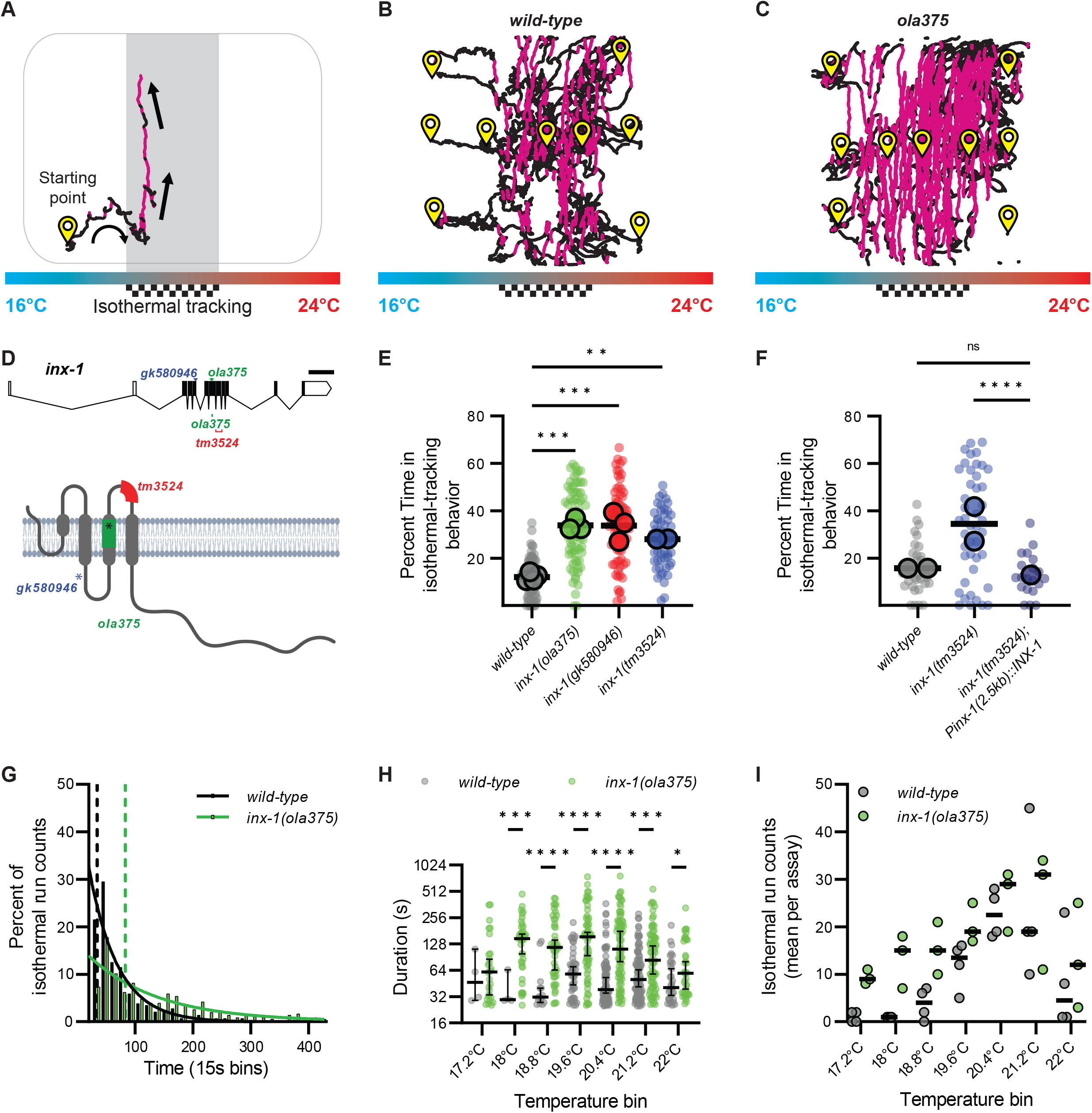
*inx-1* mutants track isotherms at context irrelevant temperatures. **A.** *C. elegans* track of a worm performing thermotaxis behavior. *C. elegans* perform two behavioral strategies during thermotaxis: gradient migration towards their preferred temperature and isothermal tracking at their preferred temperature. In the schematic, the preferred temperature region where animals are known to perform isothermal tracking is shaded and highlighted with a checkered goal pattern. Periods of isothermal tracking automatically recognized via a quantitative algorithm (see Methods) are highlighted in red. Arrows denote direction of travel. **B.** Representative image of wild-type worm tracks for animals trained at 20°C (checkered goal pattern). Animals start points denoted with yellow symbol. Animals were placed in an H-shape configuration in the gradient, as explained in Supplementary Figure 1 and Methods. Periods of isothermal tracking automatically recognized via a quantitative algorithm are highlighted in red **C.** As B, but for *ola375* mutant worms isolated from a forward genetic screen. **D.** Molecular lesions present in *inx-1* alleles, and their effects on the *inx-1* gene and protein sequenc. The schematic uses *inx-1a.1* isoform. Single nucleotide polymorphisms include *A>G* at position *X:6,948,431* for *inx-1(ola375) X*; *C>T* at position *X:6,949,062* for *inx-1(gk580946) X*). Insertion/deletions include a 16bp deletion at position *X:6,948,406..6,948,421* for *inx-1(ola375) X* and 238bp deletion at position *X:6,948,032..6,948,269* for *inx-1(tm3524) X*. Introduction of an early STOP codon in W127Opal for *inx-1(gk580946) X,* Y221Opal for *inx-1(ola375) X*. **E.** Percentage of time animals spend tracking isotherms (per worm track) for wild-type, *ola375* mutants, and two independent *inx-1* mutant alleles (*inx-1(tm3524) X* and *inx-1(gk580946) X*). Individual track values are presented by semi-transparent single-colored dots, while assay means are represented by bigger-size, slightly transparent circles with a black border. Colors denote genotypes. ** denotes P<0.005 and *** denotes P<0.0005 by Tukey’s multiple comparisons test after obtaining significance (P<0.0001) in a nested one-way ANOVA test. **F.** Percentage of time animals spend tracking isotherms, per worm track, for wild-type, *inx-1(tm3524) X* mutants, and *inx-1(tm3524) X; olaEx2136* (inx-1 rescue). **** denotes P<0.0001 by Dunnett’s T3 multiple comparisons test after obtaining significance in both Brown-Forsythe (P<0.0001) and Welch’s (P<0.0001) ANOVA tests on the individual tracks. Individual track values are presented by semi-transparent single-colored dots, while assay means are represeted with bigger-size, slightly transparent dots with a black borders. Colors denote genotypes. **G.** Histogram of the durations of wild-type (n = 288) and *inx-1(ola375) X* (n = 354) isothermal runs. Solid lines denote best-fit for one phase decay curves. Half-lives of the best-fit for one phase decay curves are 34.39 seconds for wild-type and 83.23 seconds for *inx-1(ola375) X* animals, denoted by the two vertical dotted lines. The time constants of the best-fit one phase decay curves are τ = 49.61 seconds for wild-type and τ = 120.1 seconds for *inx-1(ola375) X* animals, respectively. **H.** Semi-logarithmic (Y axis in log_2_) bee-swarm plot of isotherm-oriented run durations for wild-type and *inx-1(ola375) X,* per 0.5°C temperature bin. * Denotes q<0.05, *** denotes q<0.001, **** denotes q<0.00001 by multiple Mann-Whitney tests with a False Discovery Rate of 1% (using the Benjamini, Krieger, and Yekutieli method). **I.** Assay means of number of isotherm-oriented runs for wild-type and *inx-1(ola375) X* animals, per 0.5°C temperature bin.

To identify molecules that underlie behavioral choice, we performed an unbiased forward-genetic screen. We selected mutants that moved toward the preferred temperature but displayed defects in deploying the context-dependent isothermal tracking strategy. From this screen we isolated the mutant *ola375*, which outperformed *wild-type* animals in isothermal tracking both within and outside the ±2°C range of their preferred temperature (20°C), at the expense of gradient migration performance (example tracks in Fig. 1C, quantified in Fig. 1E). *ola375* mutant animals spent ∼34% of their time tracking isotherms, almost three times more than their wild-type counterparts (Fig. 1E). Moreover, the average run duration in the isothermal track for *ola375* mutant animals was 140.5 seconds, more than doubling the wild-type average. The distribution of their run time durations in the isothermal direction still followed an exponential decay (Fig. 1G), but the decay was two times slower than that of wild-type, with a time constant of 120.1 seconds and a half-life of 83.23 seconds. We observed that the isotherm-oriented distributions of run durations were consistently higher in *ola375* mutant animals as compared to wild type animals across the gradient (Fig. 1H), with differences being more significant near the preferred temperatures. The number of isotherm-oriented runs initiated, both for wild-type and *ola375* mutants, exhibited modulation based on the distance to their preferred temperature (Fig. 1I). Together, our data indicate that *ola375* corresponds to an allele that displays increased persistence of run duration in isotherm-oriented runs as compared to wild type animals.

To identify the genetic lesion resulting in the behavioral defects of *ola375* animals, we performed positional mapping and whole-genome sequencing (*32–35*). These strategies revealed a missense mutation and a small insertion-deletion, resulting in an early STOP codon in the fifth coding exon of the gene encoding for *INX-1/Innexin 1* (Fig. 1D and Supp. Fig 1B). Three additional lines of evidence support that *ola375* is a loss-of-function allele of *INX-1*: 1) *inx-1(tm3524)* and *inx-1(gk580946)* alleles, both loss-of-function alleles, phenocopied the *ola375* allele in the behavioral defects during thermotaxis (Fig. 1E); 2) *inx-1 (tm3524)* failed to complement the *ola375* allele (data not shown), suggesting that the *tm3524* and *ola375* alleles correspond to genetic lesions within the same gene, *inx-1*; and 3) transgenic expression of *wild-type inx-1* genomic DNA rescued the thermotaxis behavioral phenotype of *inx-1(tm3524)* mutants (Fig. 1F). INX-1 is a member of the innexin family of proteins, which are functionally and topologically related to vertebrate connexins (*36–41*). While connexins can form gap junctions in vertebrates, innexins do so in invertebrates (*37, 38, 42–49*). In *C. elegans, inx-1* is expressed in neurons and body wall muscle (*50, 51*). It contributes to electrical coupling of body wall muscle cells (*52*) and synchrony of neuronal activities during rhythmic behavior (*53, 54*).

Rescue experiments with *inx-1* cDNA using different lengths of the *inx-1* promoter revealed that expression of *wild-type inx-1* cDNA in *inx-1(tm3524)* mutants under the control of a 2.5-kb, but not a 1.5-kb promoter sequence (upstream of the *inx-1* translation initiation site) could rescue the mutant behavior. To identify the neurons where INX-1 acts to regulate the thermotaxis behavior strategies, we expressed GFP under the control of the *Pinx-1(2.5 kb)* promoter fragment (Fig. 2A) and mCherry under the *Pinx-1(1.5 kb)* (Fig. 2B) fragment, respectively, and used a subtractive strategy to identify neurons in which *inx-1* is required for rescue (Fig. 2C and Supp. Fig. 1C). This strategy led to the identification of four neuronal pairs (AIY, RIM, RIG, and an unidentified amphid neuron) that were detected with the longer (rescuing), but not the shorter *Pinx-1* promoter fragment, consistent with the hypothesis that expression of INX-1 in (some or all) of these four neuronal pairs is necessary for rescue. To further examine this hypothesis, we then generated a conditional knockout strain by flanking the *inx-1* gene with LoxP sites (Supp. Fig. 1E and (*55*)) and expressing Cre (*56*) in the candidate neurons by using specific promoters (*Pttx-3* for AIY, *Pcex-1* for RIM, and *Pceh-16* for RIG). We observed that cell-specific knockout of *inx-1* in AIY (but not in other neurons) recapitulated the aberrant action selection phenotype observed in *inx-1* mutant animals (Fig. 2E). Consistently, AIY-specific expression of wild-type *inx-1* abrogated the isothermal tracking phenotype of the *inx-1(tm3524)* mutants. The expression of wild-type *inx-1* in AIY also caused an abnormal gradient migration phenotype, presumably from *inx-1* overexpression (data not shown and (*57*)).

**Figure 2.**
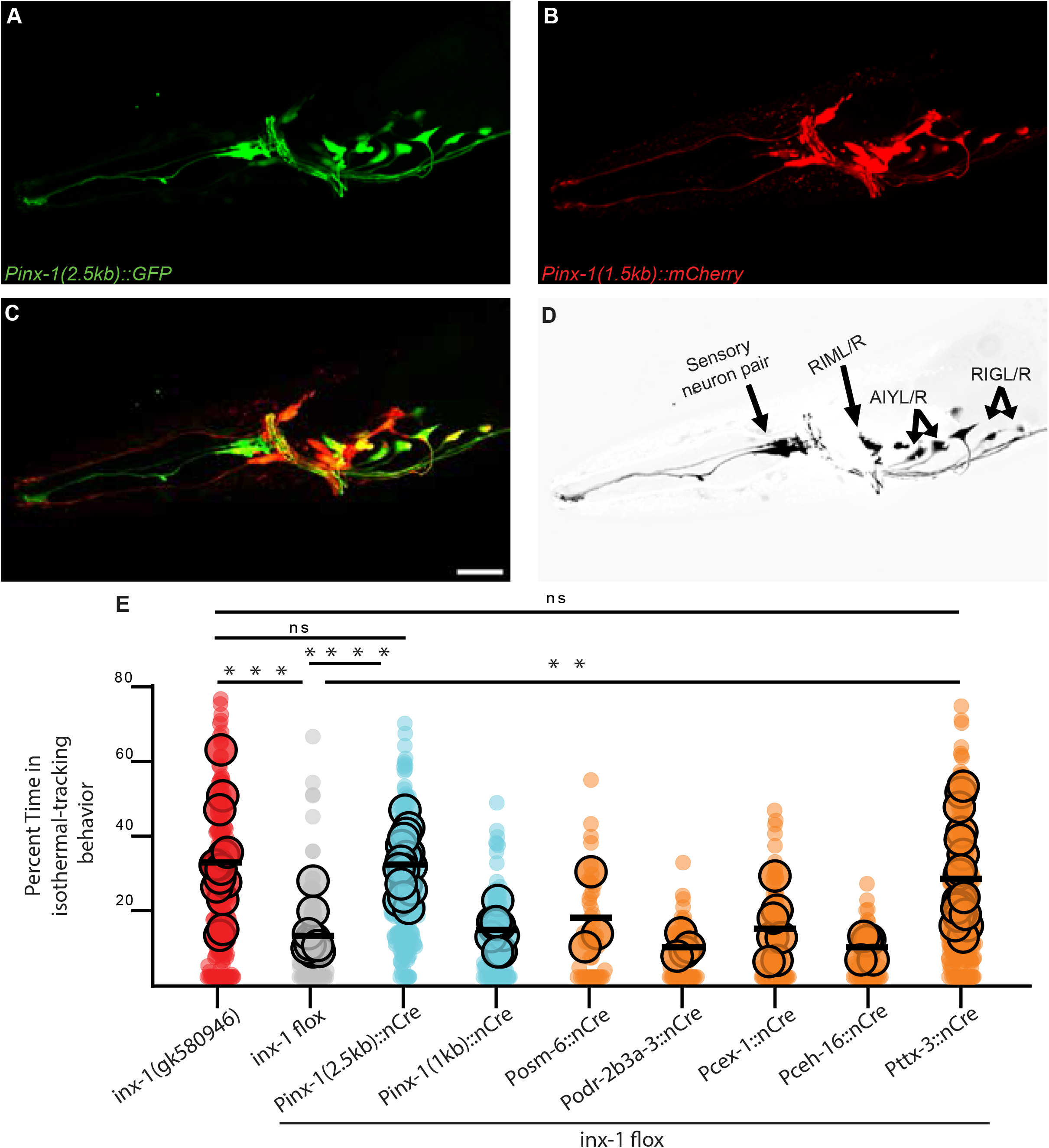
*INX-1* is required in AIY interneurons to suppress context-irrelevant isothermal tracking. **A.** Fluorescent micrograph of the head of an animal expressing GFP under the control of the rescuing 2.5kb *inx-1* promoter. **B.** Fluorescent micrograph of the head of an animal expressing mCherry under the control of the 1.5kb *inx-1* promoter. **C.** Composite of panels A-B. Scale bar is 50μm and applied to A, B and C. **D.** Neuronal pairs present under the 2.5kb *inx-1* promoter but not the 1.5kb *inx-1* promoter (for strategy, see Supplementary Figure 1C). **E.** Percentage of time animals spend tracking isotherms, per worm track, for *inx-1(gk580946)* or for *inx-1(ola278)*. *inx-1(ola278)* is an engineered *inx-1* floxed allele for conditional knockdowns using Cre recombinase (see Supplementary Figure 1E). Cre recombinase was expressed in the *inx-1* floxed allele under the indicated promoters. ** denotes P<0.01, *** denotes P<0.001, **** denotes P<0.0001 by Kruskal-Wallis test. Individual track values are presented by semi-transparent single-colored dots, while assay means are represented by bigger-size, slightly transparent dots with a black border.

The AIY neuron class consists of two bilaterally symmetric interneurons that are necessary for proper thermal gradient migration and for isothermal tracking (*11, 20, 21, 28, 30, 31, 58, 59*). They are the only known postsynaptic partners to the bilateral pair of thermosensory neurons, AFDs ((*60, 61*) and Supp. Fig. 1D). Electron microscopy studies of the *C. elegans* connectomes have predicted a putative electrical synapse between the two AIYs at their synaptic regions (*60*), but the physiological function and molecular compositions of these structures are unknown. Since INX-1 is a gap junction protein, the identification of AIYs as the site of INX-1 function promoted us to investigate whether the AIY pair is electrically coupled, and whether this coupling is dependent on INX-1. To address this, we used transgenic animals expressing the genetically-encoded calcium indicator GCaMP6 in AIY and tested the effect of depolarizing one AIY (from -60 mV to +40 mV, for 20 seconds) on the calcium dynamics of both AIYs (Fig 3A). We analyzed the calcium dynamics at the AIY synaptic terminals known as Zone 2 (*62*), where the two AIYs have been shown to respond (*24, 57*) and where electrical synapses identified by EM studies were previously reported (*60*).

**Figure 3.**
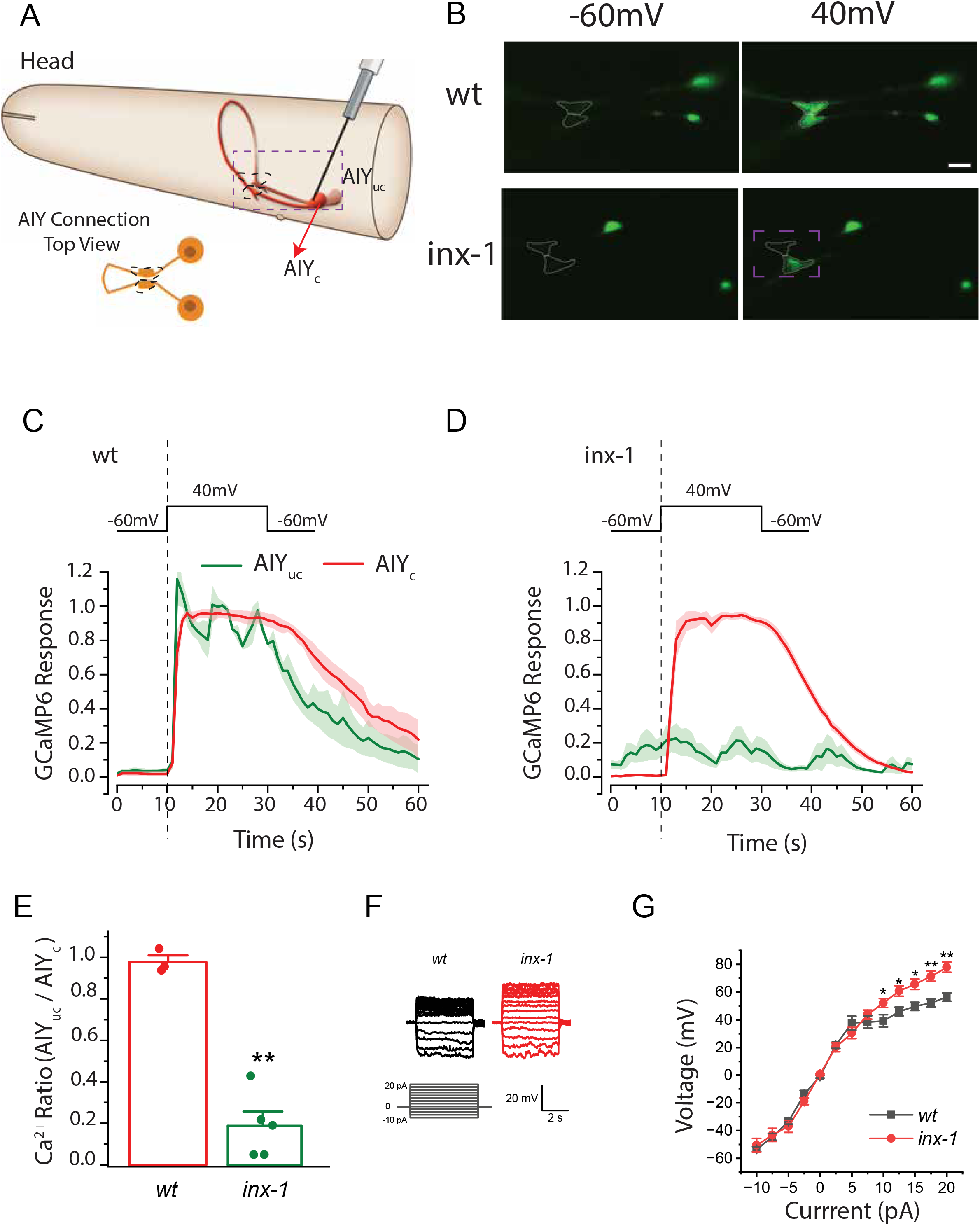
The bilateral pair of AIY interneurons are electrically coupled by INX-1 gap junctions. **A.** Schematic of the *C. elegans* head and the bilaterally symmetric pair of AIY interneurons, with the clamped AIY (AIY_c_) and unclamped AIY (AIY_uc_). Dashed box represents imaged region in B. Images in B are from dorso-ventral views of the AIY pairs, schematized in lower left cartoon, with the synaptic region (called Zone 2) highlighted with dashed lines (as also in seen in B). **B.** Sample images of GCaMP6 fluorescence in the two AIYs before and during the 40-mV voltage step in wild type (wt) and *inx-1(gk580946)* mutants. Synaptic region (called Zone 2) of the two AIYs are marked by dotted lines (also in cartoon in A). Cell bodies are also visible to the right of the synaptic region. **C.** GCaMP6 signal strength over time in clamped AIY (AIY_c_) and unclamped AIY (AIY_uc_) of wild type animals. For results of individual animals normalized by the peak fluorescent signal of AIY_c_, see Supplementary Figure 2. **D.** As in (C), but for *inx-1(gk580946)* mutant animals. **E.** Comparison of the ratio (AIY_uc_/AIY_c_) of GCaMP6 signal. The ratio is the difference between the averages of AIY_uc_ and AIY_c_ over the 20 sec depolarizing period, and the preceding 10 sec hyperpolarizing period. *wt* (*n* = 3 animals) and *inx-1* mutant (*n* = 5 animals). **F.** Sample membrane voltage traces in response to current injections for the indicated genotypes. **G.** Voltage versus current relationships of wild type (*wt*) and *inx-1* mutants (*n* = 7 animals in both groups). The averaged membrane voltage over the last 4 sec of each current injection step (5 sec in duration) was used for quantification. The asterisks * and ** indicate statistically significant differences at *p* < 0.05 and *p* < 0.01, respectively (unpaired *t*-test).

In response to the voltage step, calcium signals increased in both AIYs of wild-type animals (Fig 3B-C and Supp Fig 2). In contrast, in *inx-1(gk580946)* mutant animals, only the clamped AIY responded (Fig. 3B, D, Supp Fig. 2 and Supplementary Movies 1-2). The calcium signal ratio of the AIY pair (unclamped over clamped) during the depolarizing voltage step (+40 mV) was 0.977 ± 0.032 in wild-type and 0.187 ± 0.070 in *inx-1* mutants (Fig. 3E), indicating that INX-1 is required for the activation of the unclamped AIY. Calcium signal remained quiet prior to the voltage step in both AIYs of wild-type animals. In contrast, calcium signal often oscillated in the unclamped AIY of the *inx-1(gk580946)* mutant, and the oscillations appeared to be unrelated to the membrane voltage of the clamped AIY (Fig. 3F). Our data indicate that the hyperpolarizing voltage (-60 mV) could effectively silence the calcium activity of both AIYs in wild-type worms but not in *inx-1* mutants. Importantly, our data indicate the AIY pair is electrically coupled via INX-1.

To then determine the effect of current injections on the membrane voltage of the AIY neurons, we performed current-clamp experiments on single AIYs. In *wild-type*, the relationship between current and membrane voltage was linear over the current range from -10 to +5 pA, but exhibited a reduced slope at larger positive currents (Fig. 3G). In *inx-1(gk580946)* mutants, the slope of the membrane voltage versus current relationship was identical to that in *wild-type* at the -10pA to +5 pA range, but was substantially steeper than *wild-type* AIYs at +5 pA to +20 pA range (Fig. 3G). These findings suggest that the changes in membrane permeability of the AIYs of the *inx-1* mutants are different from those of *wild-type* animals, resulting in greater changes in membrane voltage at the current range of +5 pA to +20 pA. Collectively, these results indicate that the two AIYs are electrically coupled by gap junctions containing INX-1, and that this coupling might alter the electrophysiological properties of the two AIY neurons.

The gap junctions could serve to dampen the response of AIY to sensory inputs by shunting excitatory currents, similar to how amacrine cells in the retina are coupled via electrical synapses to achieve noise reduction during light sensory processing (*63–65*). But the observed phenotypes in *inx-1* mutants could also be influenced by uncharacterized interactions with other innexins in neighboring cells, or by undetermined *inx-1* signaling roles in AIY. To examine if the observed action selection phenotype emerged due to its specific role in electrically coupling the two AIY interneurons, we used heterologous expression of mammalian Connexin 36 (*Cx36*) and specifically expressed it in AIYs of *inx-1(tm3524)* animals. Transgenic animals expressing *Cx36* specifically in the AIY interneurons exhibited a dramatic decrease in the time spent on isothermal tracking compared to the original *inx-1(tm3524)* mutants (Fig. 4A). These results suggest that a loss of electrical coupling between the two AIYs underlies the aberrant thermotaxis behavior of the *inx-1* mutants, and that *wild-type* thermotaxis behaviors rely on the electrical coupling of the AIY interneurons via INX-1 gap junctions.

**Figure 4.**
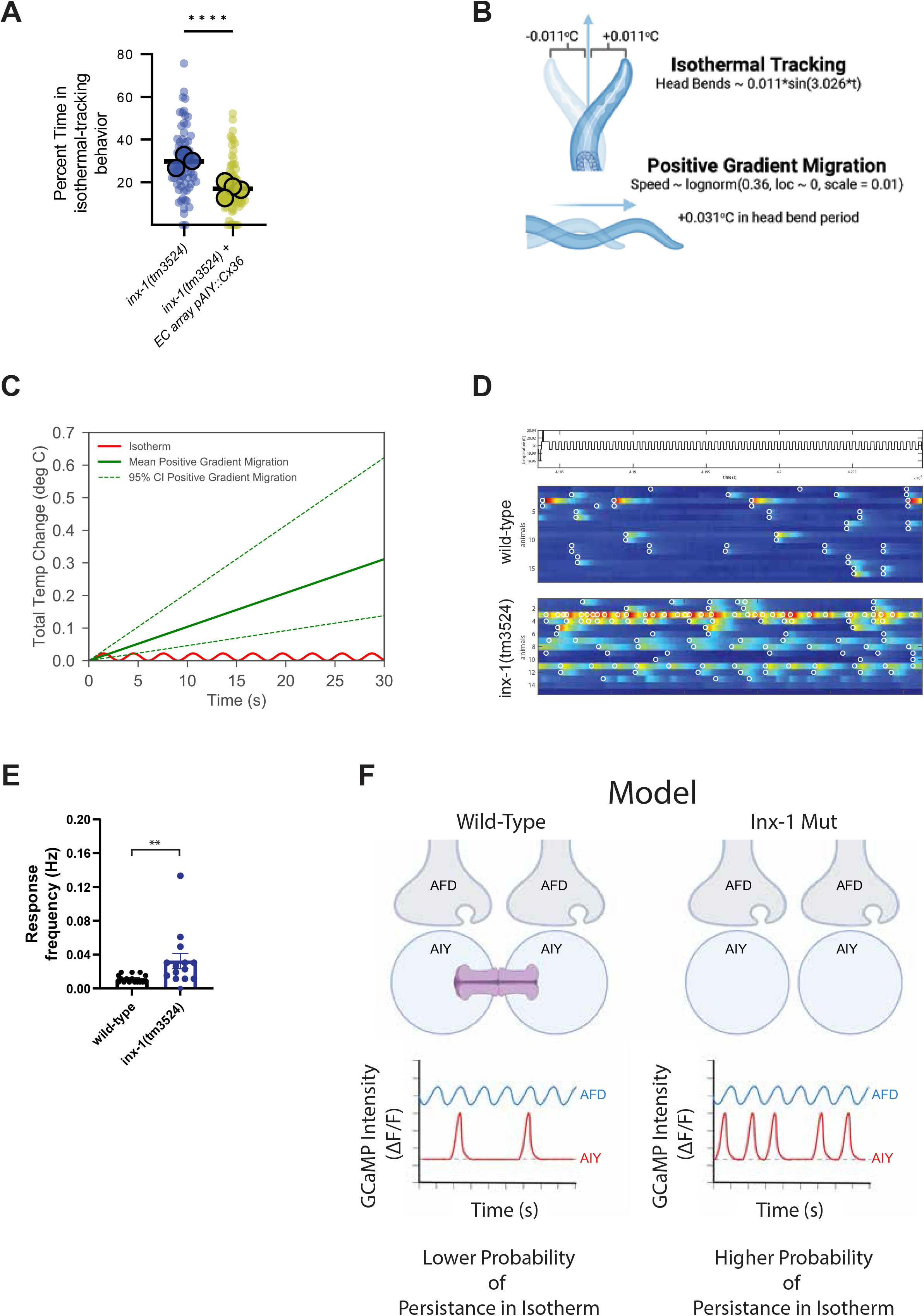
AIY sensitization in *inx-1* mutant animals increases response frequency to small temperature changes. **A.** Connexin36 was expressed in AIYs of *inx-1(tm3524)* under the control an AIY-specific promoter (*Pttx-3*), and percent time isothermal tracking was calculated for *inx-1(tm3524)* animals, or for *inx-1(tm3524)* animals expressing the *Pttx-3::connexin36* transgene. Each small circle represents a single track, and large circles represent assays. Boxes represent median and quartiles, and whiskers represent minimum and maximum points. Results of the transgenic group were obtained for three independent lines. ** Denotes *p* < 0.005 (nested t test). **B.** Diagram depicting modeled temperature changes induced by head bends in isothermal run (top) and forward movement directly up temperature gradient (bottom). **C**. Quantification of total temperature changes evoked by models presented in D as a function of run duration. **D**. Calcium responses of wild-type (middle) and *inx-1* mutants (bottom) immobilized animals when stimulated by an isotherm (+/-0.01°C oscillations around Tc 20°C, schematic on top of plots). Color scale indicates delta F/F GCaMP intensity. Responses crossing threshold are circled in white. **E.** Frequency of individual AIY calcium transients in wild-type animals (8 animals, 16 AIYs) and *inx-1(tm3524)* mutants (7 animals, 14 AIYs). Values are shown as mean ± SE and the asterisks ** denote *p* < 0.005 by two-tailed Mann-Whitney test. **F.** Schematic model of AFD to AIY signaling, and resulting behavior, in wild type versus *inx-1* mutants. In wild type animals, coupled AIYs have lower resistance and dampened responses to thermosensory stimuli coming from AFD. These dampened responses enable AIYs to integrate larger changes of thermosensory information as animals perform gradient migration. In *inx-1* mutants, uncoupled AIYs are hyperexcitable due to a change in their electrophysiological properties resulting from the uncoupling. This hyperexcitability results in AIYs responding to subthreshold sensory signals. Activation of AIYs AIY initiate and sustain a forward-moving run (*66–71*), and their hyperactivation to subthreshold stimuli would result in a higher probability of animals abnormally persisting in isotherms.

We hypothesized that the increased sensitivity of AIY observed in the *inx-1* mutants may preferentially increase the response rate of AIY to the small-scale changes in temperatures associated with isothermal tracking. To test this hypothesis, we modeled (1) head-bends by fitting a sinusoidal function to positional measurements of the nose of an animal as it freely navigates a temperature gradient (Fig. 4B and Supp. Fig. 3A), and (2) bouts of forward movement by recording the speed of animals as they move directly up a temperature gradient and fitting the data with a lognormal distribution (Fig. 4C and Supp. Fig. 3B). Our estimates suggest that animals performing isothermal tracking in a gradient similar to our experimental conditions (described in Methods and in Fig. 1A) experience oscillations, with each head bend, of + and – 0.011°C around the absolute temperature being tracked, while in the same head bend period of 3.026 seconds, animals moving up the gradient (at median speed) experience an increase in temperature of 0.031°C (Fig. 4B). We hypothesized that the smaller temperature changes experienced during isothermal tracking might result in a lower probability of activation of AIY in wild type animals, but in a higher probability of activation in the uncoupled and sensitized AIYs of *inx-1* mutant animals.

To test this hypothesis, we imaged calcium dynamics of both AIYs in immobilized wild-type and *inx-1* mutant animals after conditioning them at 20°C for several hours to create an internal state favoring isothermal tracking, and presenting them with oscillating temperature stimuli (every ∼4 seconds) centered around 20°C (with an amplitude of ± 0.01°C) (Fig 4D). In wild-type, calcium transients occurred at a low rate (0.01145 Hz, or one response every ∼87 seconds, after the animal had experienced ∼22 stimuli). These responses in wild type were often concurrent in the two AIYs, with a Pearson’s correlation coefficient of 0.8 (Fig. 4D-E). In contrast, calcium transients occurred twice more frequently in *inx-1(tm3524)* mutant animals (0.03244 Hz, or one response every ∼31 seconds) but they were asynchronous between the two AIYs (Pearson’s correlation coefficient of 0.4) (Fig. 4D-E). These results indicate that INX-1 plays a major role in the synchronous responses between the two AIYs. Our findings also support a model in which the uncoupled AIYs in *inx-1* mutants are sensitized, and respond more frequently than wild type animals to small magnitude changes in temperature (±0.02°C), like those seen during isothermal tracking.

AIY activity is known to suppress turns to induce and sustain bouts of forward movement (‘runs’) (*66–71*). In *inx-1* mutant animals, uncoupled AIYs display increased excitability and abnormally respond to small changes in temperature. Our data therefore supports a model in which AIY hyperexcitability ‘traps’ animals in long runs of minor temperature changes, decreasing the efficiency with which they move up the temperature gradient and can perform context-relevant behaviors (Fig 4F).

## Discussion

We uncovered a specific *in vivo* behavioral role played by a configuration of electrical synapses formed between bilaterally-symmetric interneurons. To differentiate between two behavioral strategies, these electrical synapses increase the effective membrane permeability of this neuronal pair to filter out sensory information of subthreshold magnitude. By coupling a pair of bilaterally symmetrical neurons, electrical synapses decrease the membrane resistance of the interneurons and dampen the effect of subthreshold excitatory synaptic inputs. This effect of electrical synapses in the circuit activity states supports context-dependent deployment of complementary behavioral strategies in the nematode *C. elegans*.

In multiple behavioral contexts, AIY calcium activity is required to initiate and sustain a forward-moving run (*66–71*). Our findings are consistent with a model in which the probability of AIY activation account for the behavioral differences observed between wild-type and *inx-1* mutants. In *wild-type* animals, AIYs have a low level of activity at the initiation of an isotherm-oriented run far away from their preferred temperature. The absence of activity would result in the probabilistic exit from the isothermal orientation via the execution of a reversal or pirouette (*31, 70*), thereby ending the run and reorienting the animal on a different direction. However, hyperexcitable AIYs in *inx-1* mutants would be activated by stimuli that are subthreshold for AIY activation of the in wild-type animals. This hyperexcitability of AIYs in *inx-1* mutants leads to the persistence of isotherm-oriented runs, and for animals to be ‘trapped’ in this behavioral state.

Wild-type animals also track isotherms, but unlike *inx-1* mutants, this behavioral strategy is restricted to a temperature context near (±2°C) of their cultivation temperature (*11, 19, 31*). We posit that *INX-1* could also serve as a regulatory switch for modulating action selection in wild-type animals. Regulation of the open/close states of the gap junctions by other molecules or post-translational modifications (*44, 47, 72–76*) may be a molecular substrate that enables a plastic uncoupling of the AIY pair. In this model, a change in the mode of sensory processing would ultimately affect action selection. Our findings also support a model whereby signal gains in the AFD→AIY synapse to smaller temperature derivatives could similarly impact isothermal tracking performance. Importantly, our findings reveal a role for electrical synapses in decreasing responses in coupled AIY interneuron pairs, which in turn are necessary to facilitates migration across the temperature gradient towards their cultivation temperature.

The organization of electrical synapses between the two AIYs might be a conserved and important configuration in sensory processing. The configuration is reminiscent of the gap junctions organization between amacrine cells in the retina. Amacrine cells are coupled by gap junctions that dampen their responses to visual sensory stimuli (*63–65*). Dampening of the gain of amacrine cells is critical for coincidence detection by photoreceptors, noise reduction and sensory processing during light adaptation (*63–65*). Thus, this configuration might confer circuits the ability to deploy context-dependent plastic responses by dynamically modulating sensory information processing, thereby increasing the versatility of neural circuits during sensory stimuli.

## Acknowledgements

We thank the members of the Colón-Ramos lab for their thoughtful comments on the project. We thank the *Caenorhabditis* Genetics Center (supported by the National Institutes of Health (NIH), Office of Research Infrastructure Programs (P40 OD010440)); the *C. elegans* Reverse Genetics Core Facility at the University of British Columbia which is part of the International *C. elegans* Gene Knockout Consortium; and Shohei Mitani (Tokyo Women’s Medical University, Tokyo, Japan) for nematode strains. We thank Bill Schafer (MRC Laboratory of Molecular Biology) for kindly providing us with their *Connexin36* construct (*77*) and Z. Altun (www.wormatlas.org) for diagrams used in figures. We thank Aravi Samuel (Harvard University) for generous assistance with technical knowledge in the development of behavioral rigs and calcium imaging platforms. We thank Hari Shroff (Janelia Research Campus) and Andrew Lauziere (University of Maryland) for image processing codes used for movies of freely moving animals. Research in the D.A.C.-R. lab was supported by NIH R01NS076558, National Science Foundation (NSF IOS 1353845), DP1NS111778 and by an HHMI Scholar Award.

## Author Contributions

A.C.C. identified the behavioral phenotype of *inx-1* mutant animals resulting in abnormal isothermal tracking and identified the AIY neurons as the site of action via a subtractive labeling strategy and the generation of a conditional Knock-Out strain for the *inx-1* gene. A.A. performed the original genetic screen that isolated the *ola375* allele. A.A.-P. identified the *ola375* allele as a genetic lesion in the *inx-1* gene, performed detailed characterization and analyses of the behavioral phenotypes associated with *inx-1* mutants, performed and analyzed calcium-imaging experiments in immobilized animals when presented with temperature stimuli, and performed and analyzed behavioral suppression experiments using orthogonal mammalian Cx36 gap junction constructs. J.B. performed the modeling experiments. L.N. and Z.W. performed and analyzed the electrophysiological experiments. I.R., E. W. and J. H. assisted in experimental design, data acquisition and analysis. A.A.-P. and D.A.C.-R. prepared the manuscript with the assistance of all authors, in particular, Z. W. and J. B. and M.G.-D.

## Materials and Methods

### Reagents and Resources

#### Molecular biology

Plasmids were generated using Gibson Assembly (New England Biolabs) or the multi-site Gateway cloning system (Invitrogen). Either Phusion or Q5 High-Fidelity DNA-polymerase (NEB) were used for cloning or subcloning elements into the Gateway entry vectors. Cell-specific promoter fragments were amplified from genomic DNA or preexisting plasmids and introduced into *pENTR41* or *pENTR 50-TOPO* vectors (Invitrogen); CDS of interest were inserted into *pDONR221[1-2]* (Invitrogen); and preexisting 3’UTR regions of commonly used genes (*unc-54, let-858*) into *pDONR221[2-3]* were used. Every insert was sequenced in their respective entry vector prior to the four-component LR recombination to generate the final expression plasmid (*78*).

#### Generation of transgenic strains

Transgenic *C. elegans* strains were generated by microinjection of the plasmids of interest into the gonad syncytia following standard approaches (*79*). Transgenic lines were selected and maintained based on the expression one or multiple of the following co-injection markers: *Punc-122::GFP, Punc-122::RFP, Punc-122::dsRed Pmyo-3::mCherry, Pelt-7::GFP::NLS* or *Pelt-7::mCherry::NLS*. Extrachromosomal arrays were integrated into the nematode genome via UV-activated trimethylpsoralen (TMP, *Sigma, T6137*), following standard methods. For a full list of strains used and generated by this work, please refer to the Supplemental Strain Table.

#### Generation of “floxed” *inx-1(ola278)* for conditional Knock-Out experiments

We inserted *LoxP* sites flanking the endogenous *inx-1* genomic coding locus via the CRISPR-based, genomic edition protocol detailed in Dickinson et al., 2015 (*55*). This strain also carries an inserted *tagRFP* sequence and a *Hygromycin B resistance* gene after the 3’ *LoxP* insertion (Supplementary Fig 1E). Complete inserted sequence can be found in Supplemental Information.

#### Nematode Strains and maintenance

Nematodes were regularly maintained at room temperature (20-23°C) or inside Precision 815 (Thermo Scientific) or I-36NL (Percival Scientific) incubators at 20°C, grown on bacterial lawns of *Escherichia coli* strain *OP50* seeded onto Nematode Growth Medium, according to husbandry standards (*80*). One-day adult hermaphrodite worms were used in all experiments unless otherwise noted. The *N2* Bristol strain was used as the wild-type background.

#### Genotyping of mutant strains

Adult worms were lysed following standard protocols and PCRs were performed using GoTaq Green Master Mix (Promega, REF-M7123). Mutant alleles were distinguished from wild-type by imaging Restriction Fragment Length Polymorphisms (RFLP) on an agarose gel or by Sanger Sequencing performed by GENEWIZ (Azenta Life Sciences). The full set of genotyping primers and PCR conditions can be found in Supplemental Information.

#### Thermotaxis Behavioral Assays

For all behavior experiments, the animals’ developmental stage was synchronized by either allowing gravid adults to lay eggs in a seeded plate for two hours, three days prior to the assay or by picking L4 animals – identified by the clear half-moon patch in the midsection of the animal – the day before the experiment. The plates were then kept in Precision 815 (Thermo Scientific) or I-36NL (Percival Scientific) incubators at 20°C up to the time of the experiment, for experiments with a cultivation temperature of 20°C, or shifted to the appropriate temperature 4-6 hours prior to testing, in the case of temperature shift assays.

Behavioral analyses were performed as described previously (*30, 57*). A population of synchronized one-day adult hermaphrodites were picked onto an unseeded plate and washed in M9 (*81*). 3-5 worms were then transferred by micropipette on a 3µl M9 droplet to the respective starting points on the assay plates (*82*), equilibrated for 5-10 min, and they were allowed to freely crawl on the arena for 30-60 min, acquiring images at 2fps with a MightEx BCE-B050-U camera. Nematode tracks were analyzed using the MagatAnalyzer software package with modifications as previously indicated (*30, 57, 83*) and additional custom MATLAB (MathWorks) scripts.

#### Sensitized forward-genetic screens

To unbiasedly find new genes that might regulate or modulate the distinct thermotaxis gradient migration and isothermal tracking behaviors, forward-genetic screens were performed on *pkc-1(nj1)* loss-of-function mutants (strain *IK105*), which perform constitutively thermophilic behaviors. This screen resulted in recovery of *ola375*. In addition to suppressing the *pkc-1(nj1)* mutant phenotype of migrating up a shallow temperature gradient regardless of their preferred trained temperature (*84*), these animals tracked isotherms more often, and further away from their preferred temperature. We mapped the causative lesion to a 5 Mb region in Chromosome X (genomic position ∼3Mb to ∼8Mb) by Hawaiian SNP mapping (*32*) and Whole-Genome Sequencing (WGS) (*33, 34*) making use of the CloudMap pipeline (*35*). Whole-Genome Sequencing (WGS) was performed by the Yale Center for Genome Analysis (YCGA).

#### Identification of *ola375* causative lesions

We further characterized the causative molecular lesion of *ola375* by fine mapping using SNPs present in the divergent, Hawaiian wild-type strain (*CB4856*) (*32*) and outcrossing SNPs with the reference N2 wild-type strain. When recombinants with wild-type DNA regions within the 5Mb previously-mapped region were recovered, from either the 3Mb or the 8Mb flank, both the suppressing *pkc-1(nj1)* phenotype and the isothermal “hypertracking” phenotype were greatly diminished. Further analysis of the behavioral phenotypes, in combination with underlying molecular lesions in that region, led us to identify a novel mutant allele for gap junction innexin gene *inx-1(ola375)*, which was exclusively responsible for the isothermal “hypertracking” phenotype under a *wild-type* background. This genetic lesion consists on both a missense SNP and a small indel in the fifth coding exon, resulting in an early STOP codon (see Fig.1D, Supplementary Figure 1 and Supp. Information). We established that *inx-1(ola375)* is the causative lesion to the isothermal tracking defects detected in this screen via four approaches: 1) examining additional alleles of *inx-*1, namely *tm3524* and *gk580946*, and determining that they phenocopy the behavioral phenotypes (persistent isothermal tracking) observed for *inx-1(ola375)*; 2) performing complementation tests to allele *tm3524* and determining that *inx-1(ola375)* fails to completement the observed behavioral phenotypes; 3) performing genetic rescue experiments with a genomic region of *inx-1* and observing that is sufficient to rescue the behavioral phenotypes and 4) performing conditional knock-out experiments and observing that cell-specific knockouts of *inx-1* in the AIY interneurons are sufficient to reconstitute the observed behavioral phenotype for *inx-1(ola375)*.

### Major Types of Behavioral Paradigms Used

#### Shallow gradients for Gradient Migration Quantification

The original suppressor screen was performed on equipment previously described (*30, 57*), monitoring nematode gradient migration in the presence of a shallow temperature gradient (0.18°C/cm). Briefly, two pairs of thermoelectric components controlled by two Accuthermo FTC100D PID controllers sit at either side of an aluminum slab, and generate a defined linear temperature gradient. The system is cooled by a closed refrigeration system connected to a liquid cooling radiator in contact with dry ice. The aluminum slab in turn contacts a square assay plate (Corning®) with a 224 × 224 mm internal arena where the worms will perform. To ensure efficient heat transfer between the slab and the arena, either a volume of glycerol was used or a fitted, smaller aluminum sheet was intercalated between the aluminum slab and the assay plate. Red LEDs parallel to the plate generate a dark background image with bright outlines of the nematodes, captured by a MightEx camera (BCE-B050-U) above, at 2 frames per second, for 30-60 min. The whole system is encased in a modified cabinet. Unless otherwise explicitly noted, the gradient of the arena goes from 18°C to 22°C, and animals are placed in the middle of the arena, near 20°C. 24-33 animals are tested per assay.

#### Moderate and steep gradients for Isothermal Tracking Quantification

*C. elegans* perform maximal isothermal tracking behavior at ∼0.6°C/cm gradients or higher (*31*). To generate these gradients, we used a modified, smaller version of the equipment described above and previously (*30, 57*), kindly gifted to us by Aravi Samuel (Harvard University). Unless otherwise explicitly noted, the gradient on the arena is centered on 20°C and goes from 17°C to 23°C for 0.6°C/cm gradient and 16°C to 24°C for 0.8°C/cm gradient. To quantitatively assess and adequately quantify isothermal tracking across the full gradient, a population of animals is assayed by starting in an H configuration, as shown in Supplementary Figure 1A. For a qualitative assessment of animals performing more isothermal tracking or under the wrong context, four starting droplets at each respective edge of the gradient were used. 24-27 animals are tested per assay.

### Imaging

#### Confocal imaging

Young adults or L4 animals were mounted in 2% agarose dissolved in M9 buffer pads and anaesthetized with 10mM levamisole (Sigma). Confocal images were acquired with dual Hamamatsu ORCA-FUSIONBT SCMOS cameras on a Nikon Ti2-E Inverted Microscope using a confocal spinning disk CSU-W1 System, 488nm and 561nm laser lines and a CFI PLAN APO LAMBDA 60X OIL objective. Images were captured using the NIS-ELEMENTS software, with 2048px × 2048px, 16-bit depth, 300nm step size, 300ms of exposure time and enough sections to cover the whole worm depth.

#### Calcium Imaging

Imaging calcium dynamics was performed as previously described (*57*), with some modifications. The sample mounting protocol was modified to enrich the samples with animals positioned dorsoventrally, allowing for imaging of both AIY neurons simultaneously. Temperature control elements and most microscopy elements remain identical to (*57*), with a Leica DM6B being used in addition of Leica DM5500. Image acquisition was performed using MicroManager (*85*).

### Quantification and statistical analysis

#### Quantification of isothermal tracking

Worm tracks were first analyzed and segmented by a modified MAGATAnalyzer software package (*30, 83*). These trajectories were then filtered into periods of isothermal tracking, defined as forward motion events in which at least 90% of the displacement occurred in the vertical, isotherm orientation, for a minimum of 25 seconds; and periods of non-isothermal tracking in which the movement of the worm did not pass the isothermal tracking filter. The segmented isothermal tracking periods were further analyzed by their duration in seconds, temperature at which the period started and number of events.

#### Quantification of behavior

Quantifications of turns, thermotaxis indices and other parameters relevant to gradient migration were automatically scored per worm track by an adapted MAGATAnalyzer software package, previously described (*30, 83*).

#### Quantification of calcium imaging in AIY

Segmentation into regions of interest and downstream data processing was performed using FIJI (*86*), and custom scripts written in MATLAB (MathWorks) as detailed previously (*57*). For analyses of AIY calcium dynamics, we generated and quantified a ROI at the synaptic subcellular Zone 2 region (*62*). Responses were scored as the initial rise of the AIY calcium signal as determined by a human observer and an automated response calling based on signal intensity and its derivative, as previously described (*57*).

#### Electrophysiological analyses

Electrophysiological analyses were performed with transgenic strains expressing *Pmod-1::GCaMP6s* and *Pttx-3::mCherry* in *wild-type* and *inx-1(gk580946)* mutant genetic backgrounds. In each experiment, a young adult hermaphrodite animal was immobilized on a Sylgard-coated circular coverglass by applying Vetbond Tissue Adhesive (3M Company, St. Paul, MN) along the anterior dorsal region. After a longitudinal cut (∼ 200 µm) was made by a diamond dissecting tool in the glued area, the cuticle above the cut line was pulled back and glued onto the coverglass to expose head neurons. The coverglass was then transferred to a recording chamber containing the extracellular solution, which contained (in mM) NaCl 140, KCl 5, CaCl_2_ 5, MgCl2 5, dextrose 11 and HEPES 5 (pH 7.2). Following identification of the two AIYs based on mCherry fluorescence, one of them was used for voltage- or current-clamp recording in the classical whole-cell configuration. In the voltage-clamp experiments, AIY was held at -60 mV and stepped to 40 mV for 20 seconds before returning to the holding voltage. Meanwhile, calcium transients of both AIYs before (10 sec), during (20 sec), and after (30 sec) the voltage step were imaged at 1-sec intervals using an electron-multiplying CCD camera (iXonEMþ885, Andor Technology, Belfast, Northern Ireland), a FITC filter set (59222, Chroma Technology Corp.), a light source (Lambda XL, Sutter Instrument), and the NIS-Elements software (Nikon). TTL signals from the camera were used to synchronize the recordings of calcium transients with the voltage-clamp protocol. In the current-clamp experiments, negative and positive currents over the range of -10 pA to +20 pA at 2.5-pA intervals were injected into the clamped AIY for 5 sec per step. Borosilicate glass pipettes with a tip resistance of 3B5MO were used as electrodes for current- and voltage-clamp recordings with a Multiclamp 700B amplifier (Molecular Devices, Sunnyvale, CA), a digitizer (Digidata 1440A, Molecular Devices), and the Clampex software (version 11, Molecular Devices). Data were sampled at a rate of 10 kHz after filtering at 2 kHz. The pipette solution contained (in mM) KCl 120, KOH 20, Tris 5, CaCl2 0.25, MgCl2 4, sucrose 36, EGTA 5 and Na2ATP 4 (pH 7.2). A Nikon FN-1 microscope equipped with a 40X water-immersion objective was used in the electrophysiological and calcium imaging experiments.

### Statistical analyses

All statistical tests were performed using GraphPad Prism version 9 for Windows, GraphPad Software, San Diego, California USA, www.graphpad.com Chosen statistical tests are described in the relevant figure legends.

#### Supplementary Figure Legends

**Supplementary Figure 1. Strategies, strains and constructs** related to Figure 1. **A.** Schematic of thermotaxis assays and behavior. (Left) Schematic of the behavioral choice assays used in most studies, in which animals are placed at the middle of the gradient, and allowed to migrate towards their preferred temperature region (shaded grey), where they perform isothermal tracking. (Right) Schematic of the assay using in this study, in which animal start sites (circles) were placed in an “H” configuration and trained to prefer 20C (shaded area in middle of assay), as to better capture behaviors of animals performing gradient migration up or down the gradient as they transition into isothermal tracking. **B.** Schematic and sequence information for the *inx-1(ola375)* allele isolated in this study from forward genetic screens. **C.** Schematic of the substractive labeling strategy to identify the INX-1 site of action. A promoter fragment of 2.5kb can drive expression of *inx-1* cDNA and rescue the observed thermotaxis defects for the *inx-1* mutants, while a promoter fragment of 1.5 kb is insufficient to do so. By creating transcriptional fusions of both promoter fragments, we identified candidate neurons which are uniquely labeled by the rescuing promoter fragment. **D.** Schematic of part of the thermotaxis circuit, highlighting the AIY interneuron position as the primary interneurons downstream of the thermosensory neuron AFD. **E.** Schematic of *inx-1(ola278)* a floxed allele engineered for conditional knockdowns of the *inx-1* gene.

**Supplementary Figure 2. Examination of AIY coupling by INX-1, related to Figure 3.** GCaMP6 signal strength over time in clamped AIY (AIY_c_) and unclamped AIY (AIY_uc_) of wild type (A and C) and the *inx-1* mutants (B and D). Shown here are results of individual animals normalized by the peak fluorescent signal of AIY_c_. C and D are the same as Figure 3 C and D, and represent the cumulative results of the individual animals.

**Supplementary Figure 3. Thermotaxis modeling parameterization and Pearson coefficient firing between AIY pairs. A.** Data from a freely moving animal during a straight run, displaying the position of the nose tip (dots) and fit with a sinusoidal curve. **B.** Histogram of the speeds of runs of animals trained at 25C and placed at 20C, and moving up the gradient towards their preferred temperature, with a lognormal fit (in red). **C.** Pearson’s coefficient in AIY pairs between wild-type animals (n = 8) and *inx-1(tm3524)* (n = 7). Values are shown as mean ± SE. the asterisks *** denote *p* < 0.0005 from two-tailed Mann-Whitney test.

## Supplemental Movies

**Movie 1**. Calcium responses of AIYs (ventral view) in wild type animals stimulated by simulated isotherm (+/-0.01°C oscillations surrounding Tc, 20°C, related for Figure 4D)

**Movie 2**. As Movie 1, but in *inx-1(tm3524)* mutants.

## Supplemental Strain Table

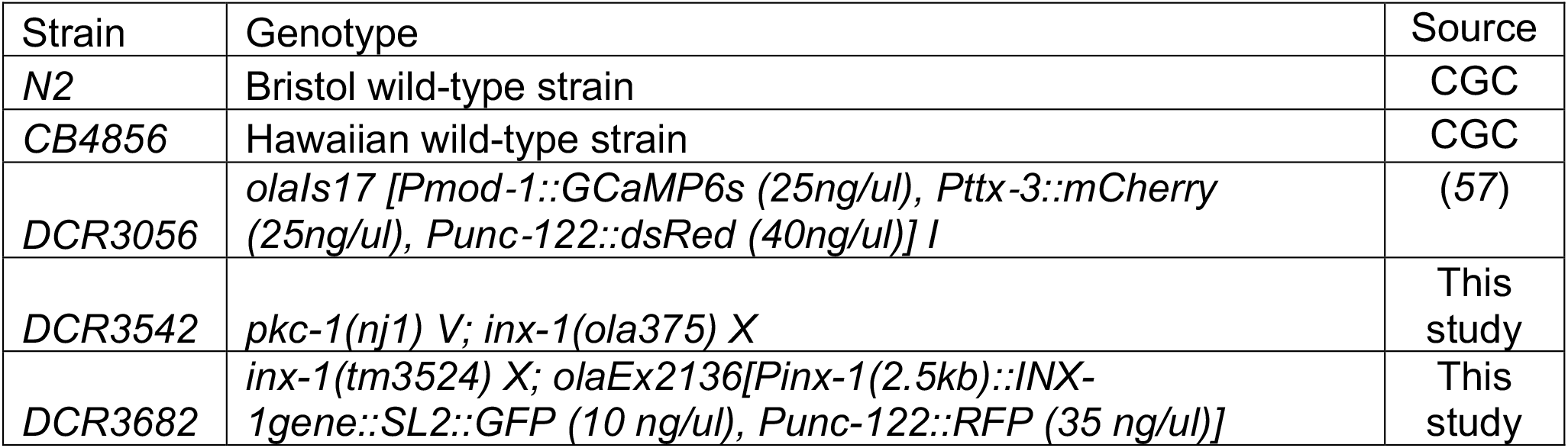
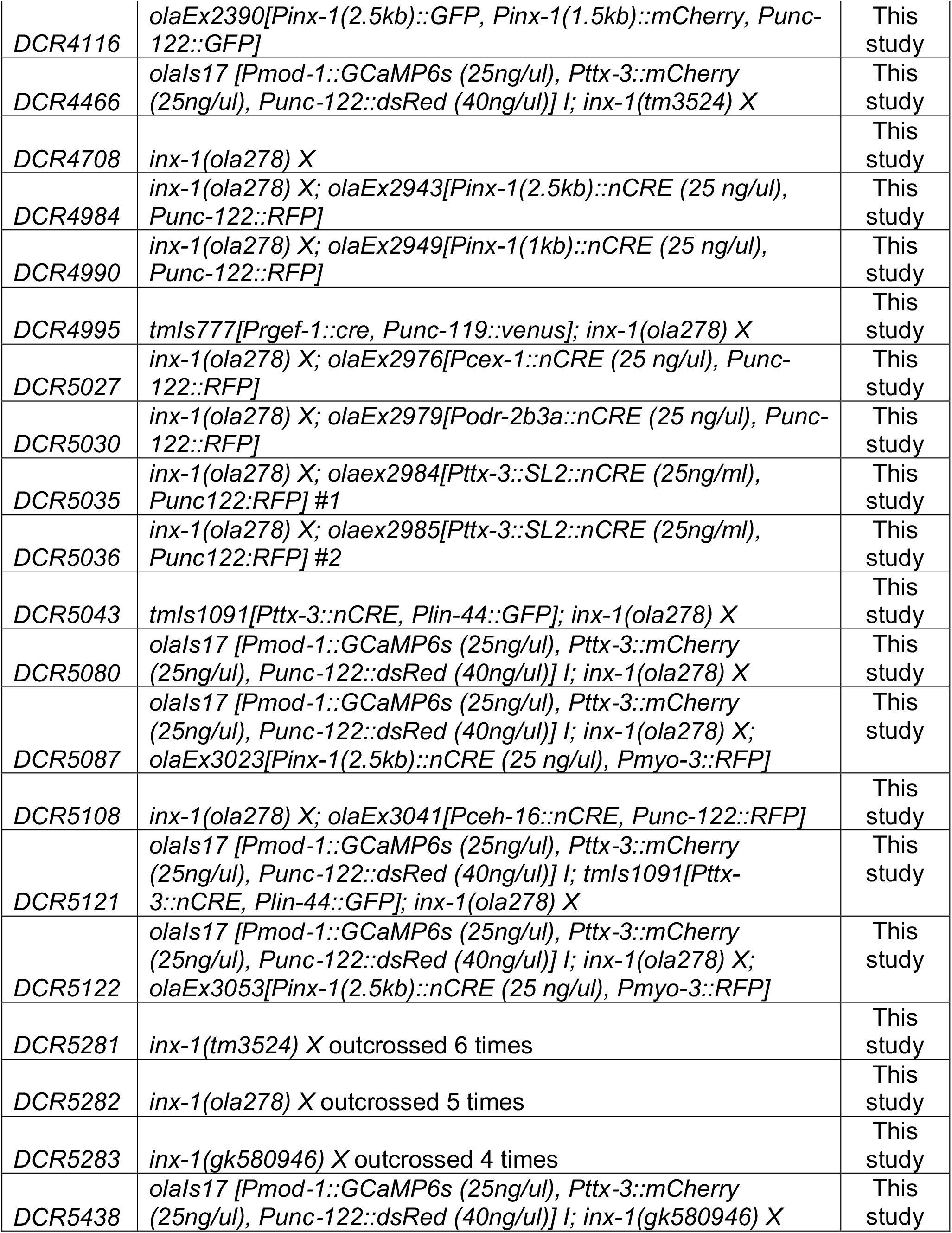
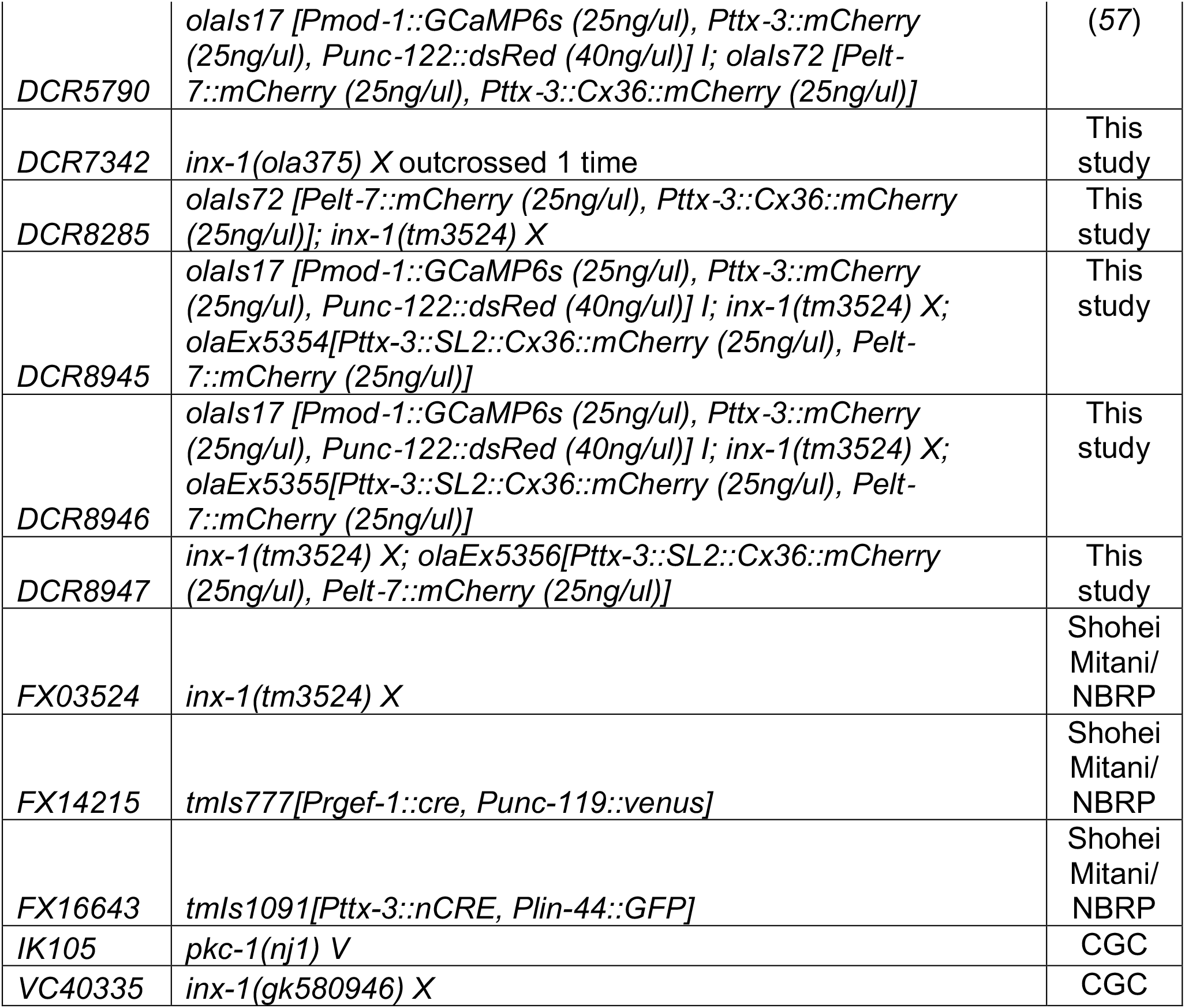

### *inx-1(ola278)* sequence (insertions in bold)

gaggcacagtttgaaaataaattaaattttaattcatttgatgattttgttttctcttgaggcttaaaaatgataaacggtacaaa actacaaaaaaactccataagtctttattttcttaatttttgaaattttatttcaaaatgcacaagatccatttatcatatttatagttct tcatcttcttttttttaatatcctctttttgtgttttattcggtccatacgtgctcttcaacttttttttcatttttgactttaaataagaaaagat ggcaataaaaaatttggccgaagagggcgatggacggatgaaaatctactaaaaggattataactcaatttgatatgctct tcgggaggtatcctcctgacgagatggaaaagaagaagaagaagaagaagaagcttgtcatcgtttcatcggcaaaga gacgggtggacattagaccacgccccacagggaaacctttcgagtttcatccacctctgtctgtttaacatatttgtttttttcta gctctatttttcttccgcttctactgcgtacttttgcataattctatttctaagtctgatcattataagtatcatcctgaacatcgcaca ctaaacatcctcggc**ATAACTTCGTATAGCATACATTATACGAAGTTAT**cggcacggatgaagta gttttcattgcagttcttgtccgccggaATGCTTCTATATTATCTGGCGGCCATATTCAAGGGCTTA CATCCGCGAGTCGACGACGATTTTGTGGACAAGCTCAATTATCACTATACTTCTGC TATTATATTCGCGTTTGCGATTATTGTGTCTGCCAAGCAGTACGTAGgtaagtctgatttcat taattcagcttctctgcgcctaccttttcacgttaaaatcaattactttcagGTTATCCGATACAATGTTGGGTG CCTGCGCAGTTCACCGATGCTTGGGAACAGTACACCGAAAACTATTGTTGGGTGG AAAACACATACTACCTCCCGTTAACAAGTGCATTTCCATTAGAATACGGTGACAGG AGgtaatttaaaacattcaagcttattcagaaacttttcttgttacagGGCACGACAAATCAGTTACTATCAA TGGGTGCCGTTTGTGTTAGCGCTCGAAGCGTTATGTTTCTACATCCCGTGCATAAT GTGGAGAGGACTGCTGCACTGGCATTCTGgtaagataatgaaacgggagatgaaaccaaagaaa aaagaacttccaactgacgatattggtcgttaaaaccgcacaactgtgcaatctgtgtgtgtctgtgtatacgtatgtgccaa ggcctgagatgatgaaaagaatgaaaaaagaaaagaaagtagtctagagaaaagtaagtgtgaagtggagaggcga tccattaataatttattgcctaacgattctagagaaacttttatcaccttggaagaaaaatgagggccttgatcatgaaagctg cgagagaaggatgtgttcggaacgcagcgtgccatttgacataagtagacgggagacctacgttgttttatttttcagGAA TCAATGTGCAATCATTGACTCAAATGGCATGCGATGCACGAATGATGGATGCTGAT GCCAGGGCAGCCACCGTGCAGACAATTGCAGGGCACATGGAAGACGCTCTTGAA ATTCAACGAGAGgtctgtggaagtcatgttggtgaataacaaaatcaactctattttcagGTCACCGATGTG TCGGGCATGTGTGTCCAGAAGCGATGGGCAAACTATGTGACATTACTATATGTATT TATTAAAATGCTCTACCTTGGAAATGTTGTATTACAAGTGTTTATGTTAAACAGTTTC CTTGGTACTGACAACCTTTTTTACGGCTTTCACATTTTGAGAGATTTGTTGAATGGT CGTGAATGGGAAGTTAGTGGGAACTTTCCACGTGTAACTATGTGTGATTTCGAGgta aaacaacgtggattagtatcagttttaaaaaatgttttaagGTACGAGTACTGGGTAATGTGCATCATCA TACAGTCCAATGCGTGCTAATGATTAACATGTTTAATGAGAAAATATTTTTGTTCCTT TGGTTCTGGTACTTCATGGTAGCTTTTGTGTCAGCAGTGTCCATGTTCCATTGGATT ATTATATCTTTCTTACCAGGACAGgtaaccgaaataccctttttttatgatttacttaacttttgtttttcattttccag CACATGAAGTTCATCAGAAAATATCTACGAGCTACAGATTTGGCAACTGACAGGCA GTCGGTGAAAAAGTTTGTTCACAAGTTCCTCGGATATGATGGAgtatgttatgattttcgaaac acttcactaacaagatgtatttagGTGTTTTGTATGAGAATGATTTCGGCACATGCTGGAGATAT TCTTGCTACAGAACTAATTGTTGCTCTGTGGCATAACTTCAATGATCGTGTCAGGAA Ggtgagctaataagatcggctaagttgcggattggtttgcttctcaataagttttcattctaatttcaacccagcaatagtattca atttttatttttcatttatttcaaacaacctgaaacaaaatcaattaaaaagacaaaaaaaaaccacgacagatcgaataag tcaaatgagtggccgacgaatccaatccctaaaggcatatcatttctattgtttccagaatttcgagcaagtctctgagcagta gacgccaaatcctccaacagcgcttctatctctctatctatcaatgttggtttttccattctaatgtagctttgtctgaaaaaaaat gcgccccgccccctggaaacgcataacagcctatcaaactcactaaccagtttatagtttttgtgtccaccatccaccgaag tgcctttagtgattgattcgattctttgcccattcaactgattattttgtctttgtttcatttttcgccgtcacacactgcactatttttttttta aaacgtttggatcccagtttccacgttgcaaaaagcaaagtggtgtaactcaagaatttgaaatttcaagaaagattatctg gattgaatataattctaatttcccttttccaattcacacttacacattcacatacacatttttccctcttctcttcctctaactcacaca gtcactcacacacactgtatcgtaatctgtgattttttttttcttccaattacagtatatcgaatataaagtgtagaagactgcatg atccgttgtagaattaatatagtcggatctgaaactaacacacaactttctatcacttttcatggctactgtagtagttctttcata gtgcctgtgtttgtccgcttgtgatgggtggtcatgtgtgaaaatcttcctaattttaaaactgtgcattctgaaaaaaagtttcga aacatttatactgaatcaaagttttgtagaacaatttttgtgaacaagtaaaaggttaacttgtacaatcatcgcaaacgaga ccactcatacaatcggataagcacaaacacgcaattttctgcaagaataattcgattgaattttttgctttccattgtaaccgatt tgaagatacaattcaactcaacggaatccctcaaccactaaccaccacaactcaccactacctccagccaacacgtatgt ccgaaccgggtttttggtttccataacagagtaacacagcgagagtcagttgtatgtagtcgtatcttttttcggaatgcactct ctttttatattttttgaacctattttgtcgacaatttgccttcttcacccatttatcaccatgatccccaccatcaggcggttggcatgc aattagattagatgtactcaccggttgtgcctggctttctgtttttcttcttgtttttttctgactttcaacaatctgtctgcacaaatttgt ttactctgaaaacctggtccgattcttgtatcgaatatgtaaacaacgatatgcctgcccctaaattcctatgtttcccgttttgaa aaattttgatataattctataatttgagaagtaaaaaagaatgcatgaaaaaaacgaatttaatataattcagtgctatattact tgagttctccaacccaagacttgcttttgtttcattagtgtactacattgattttgattccttgattttgatttgataccaaaccgatttt caaggggtggcaaaacgcaatgaatgcctgtgtgacaggcggctcctacaaatttcattagataacctaaaacgcttgcc acttacagAGTCCAATCGAGATGTTCGAAGGCGGTGTCACTCAGTCACCTAGCAAATT AGACGCCAATTTCAAAACGTGGCTTCTCGGCCAGACGAGgtgagtcaagatggacacatgta ctgacacgctgacatgttgaaaattgcaaaattagacttcagcttgtcaacgtgtttttgtttcaagtgacgtgatagtaatttgt aaatgtcaaagtatatgtatggatgtgcgtgtgtgtacggcaaaactgttagggtttgtgaacttgctttaaaaagttagatgta aatgttaaatatatttcaaagtctctttcatttgtagaaacaaactattttttattaacttaaaagcagggctgtgcggaagttaga tcaaatttgcaatgcggtttcatttgtatagatttacccctaaactagtcattggcgcaaagtatgtagaaaaaacatgcgcac aagttttgcaaatactttttagccaaccagctgactgagagcgaataaatatttttctaaattttcttgacgaagaagtgtcaca acttattcaacctattcaaattctataattgttctcaaattaatttttgcgaacacaataaaaagtttaaatatgctctagcagctc aatttaccagccatgatcagaattatcgtataaaataaattcaaagcagaataactagttgactttttattttatgaagaacata tctttattttcaaaaacattaagagctccaaaaagtaaagctcaaaaattgagaggaaattctatgcgaacttttttaacaatt caaaataaatgacaagttcacaaaccattattagtgattagctagcagacagaaaagctctttagtgttcgcaaaaatgtg aaaatttttgacaaaatgtaaaatcctagaatagttgagcgagagcgcactgattggtcgagcctgtgttttgttgtattgcag ggcggaggccaacgggggctctcggctgctgtctgctcgctaatgtggcgccgatttcgccccaagtgtcagaatcccaat catcctacaccgtgtgtcctcacttcaagttggaattgaattgcagGGGCAAGCCGCCCTTCGACGGCTCA AATCCGACGCGTGGAAAGAAACGCCGAAAATCAGATGGTTACTTCACGTTCGTCTA Atatcttgcattcaactattcaaccttcccgagttctttcctcacttctcatctaacccgaccacgtggtgctcgcctatcgtccac atctttcaattatttgataatatgtatttctgtgccatttcatgtttttattcaattttcaaaatgttttgttttattatttggtgaatttagttcg aatattccaacttttataaccaaaataattgtacaaagcattatgaaatgtgtcataagttcttctctgaatcattcaaatacctg ggtactttttcgaccccgttctgaaacaaaattgagcacagctcttgttctaccttttcccatcccccagtgccaatattttaggtg ctctctttccaagaaatctcaaacttcacacatcgacacgccataaacctttggtaaacccttacccaaatcaaaaaaaag ctgagtgaattctagtcatttgttgtacagcccctttcctccccaactatttctgaccctcttttctgacaaatgttggactatttaatt gtctttcctacaaaaaaatacctttgtgaactgcacttctcaatttacaaaacgatacagtgactgtttatagcgggtaataag gcaacgtgcagtaatccattctaataagtttaggcactataacttaatcgtatttctgatttttcttttctcatctaaatttataggaat tttcaaaatttcaaaattgttccctttcaaattctcttagggaattattttctatacaattttcatcgggtgtatggtcccctcttcccttc ttttcgcccccaacgacacaatttttttttcaaatttaggaatttttcccttttcctttttctttctattggattttcatttctttccatcggtga ctcagataacaattattaccttcataatccatcaattacaaaagaacctaacaactttccaatgcatttcacct**ATAACTT CGTATAGCATACATTATACGAAGTTATAAAATGGTGAGTGTGTCTAAGGGCGAAGAGCTGATTAAGGAGAACATGCACATGAAGCTGTACATGGAGGGCACCGTGAACAACCACCACTTCAAGTGCACATCCGAGGGCGAAGGCAAGCCCTACGAGGGCACCCAGACCATGAGAATCAAGGTGGTCGAGGGCGGgtaagtttaaacatatatatactaactaaccctgattatttaaattttcagCCCTCTCCCCTTCGCCTTCGACATCCTGGCTACCAGCTTCATGTACGGCAGCAGAACCTTCATCAACCACACCCAGGGCATCCCCGACTTCTTTAAGCAGTCCTTCCCTGAGGGCTTCACATGGGAGAGAGTCACCACATACGAAGACGGGGGCGTGCTGACCGCTACCCAGGACACCAGCCTCCAGGACGGCTGCCTCATCTACAACGTCAAGATCAGAGGGGTGAACTTCCCATCCAACGGgtaagtttaaacagttcggtactaactaaccatacatatttaaattttcagCCCTGTGATGCAGAAGAAAACACTCGGCTGGGAGGCCAACACCGAGATGCTGTACCCCGCTGACGGCGGCCTGGAAGGCAGAAcCGACATGGCCCTGAAGCTCGTGGGCGGGGGCCACCTGATCTGCAACTTCAAGACCACATACAGATCCAAGAAACCCGCTAAGAACCTCAAGATGCCCGGCGTCTACTATGTGGACCACAGACTGGAAAGAATCAAGGAGGgtaagtttaaacatgattttactaactaactaatctgatttaaattttcagCCGACAAAGAGACCTACGTCGAGCAGCACGAGGTGGCTGTGGCCAGATACTGCGACCTCCCTAGCAAACTGGGGCACAAACTTAATTCCGGAtaaGGAT GATCGACGCCaACGTCGTTGAATTTTCAAATTTTAAATACTGAATATTTGTTTTTTTTCCTATTATTTATTTATTCTCTTTGTGTTTTTTTTCTTGCTTTCTAAAAAATTAATTCAATCCAAATCTAAacatttttttttctctttccgtctcccaattcgtattccgctcctctcatctgaacacaatgtgcaagtttatttatcttctcgctttcatttcattaggacgtggggggaattggtggaagggggaaacacacaaaaggatgatggaaatgaaataaggacacacaatatgcaacaacattcaattcagaaatatggaggaaggtttaaaagaaaacataaaaatatatagaggaggaaggaaaactagtaaaaaataagcaaagaaattaggcgaacgatgAGAATTGTCCTCGCTTGGATTTTTGCTTTCGTCGTAAATCTACACACGCGTCTCTTCCGTGCGAGAGTCCAAGCCAGCAGCCAAATTCGTTGACTGAGTATTCAACGTTTATACGTTGTCGGCAACGAGAAATAGGAAAATGCATCGGGAAATGTTCTTTTTTCGATTTTTTCCAAGGTTTTGACAAATTTTACCACGAATTTTGCTATGTTTTCAATTAAAAAATATGTTATTCAACTGTTTCTATGAGGAAAATAAGGCTTTGCATGTAATTTTCTTATTCAGCATAATTTTTAATTAATTTGAATTTTCTGTCCTAACGTTTATTTTGTTTTCTTGGTTATGACTGATCTGAAATTAATTTTTGAATTTTAAGGTAATATGTCAGGCGGTGCCGCAAGTTTGTACAAAAAAGCAGGCTCCATGAAAAAGCCTGAACTCACCGCGACGTCTGTCGAGAAGTTTCTGATCGAAAAGTTCGACAGCGTCTCCGACCTGATGCAGCTCTCGGAGGGCGAAGAATCTCGTGCTTTCAGCTTCGATGTAGGAGGGCGTGGATATGTCCTGCGGGTAAATAGCTGCGCCGATGGTTTCTACAAAGATCGTTATGTTTATCGGCACTTTGCATCGGCCGCGCTCCCGATTCCGGAAGTGCTTGACATTGGGGAATTCAGCGAGAGCCTGACCTATTGCATCTCCCGCCGTGCACAGGGTGTCACGTTGCAAGACCTGCCTGAAACCGAACTGCCCGCTGTTCTGCAGCCGGTCGCGGAGGCCATGGATGCGATCGCTGCGGCCGATCTTAGCCAGACGAGCGGGTTCGGCCCATTCGGACCGCAAGGAATCGGTCAATACACTACATGGCGTGATTTCATATGCGCGATTGCTGATCCCCATGTGTATCACTGGCAAACTGTGATGGACGACACCGTCAGTGCGTCCGTCGCGCAGGCTCTCGATGAGCTGATGCTTTGGGCCGAGGACTGCCCCGAAGTCCGGCACCTCGTGCACGCGGATTTCGGCTCCAACAATGTCCTGACGGACAATGGCCGCATAACAGCGGTCATTGACTGGAGCGAGGCGATGTTCGGGGATTCCCAATACGAGGTCGCCAACATCTTCTTCTGGAGGCCGTGGTTGGCTTGTATGGAGCAGCAGACGCGCTACTTCGAGCGGAGGCATCCGGAGCTTGCAGGATCGCCGCGGCTCCGGGCGTATATGCTCCGCATTGGTCTTGACCAACTCTATCAGAGCTTGGTTGACGGCAATTTCGATGATGCAGCTTGGGCGCAGGGTCGATGCGACGCAATCGTCCGATCCGGAGCCGGGACTGTCGGGCGTACACAAATCGCCCGCAGAAGCGCGGCCGTCTGGACCGATGGCTGTGTAGAAGTACTCGCCGATAGTGGAAACCGACGCCCCAGCACTCGTCCGAGGGCAAAGGAATAGACCCAGCTTTCTTGTACAAAGTGGGTCCAATTACTCTTCAACATCCCTACATGCTCTTTCTCCCTGTGCTCCCACCCCCTATTTTTGTTATTATCAAAAAACTTCTCTTAATTTCTTTGTTTTTTAGCTTCTTTTAAGTCACCTCTAACAATGAAATTGTGTAGATTCAAAAATAGAATTAATTCGTAATAAAAAGTCGAAAAAAATTGTGCTCCCTCCCCCCATTAATAATAATTCTATCCCAAAATCTACACAATGTTCTGTGTACACTTCTTATGTTTTTTACTTCTGATAAATTTTTTTGAAACATCATAGAAAAAACCGCACACAAAATACCTTATCATATGTTACGTTTCAGTTTATGACCGC AATTTTTA**cagaggtagttttccttatttgtttggaattttgcatggtgcattttctgccttcccatttttcttgcaaaaataaaat gttctaaatttttgatgtttaaaattatttttttttcaattttaagaatattcattgattgaaaaatcataaatcaaaattctacatcaaa atatgaatttaaacattcgcctgattgaaaaaatcactgagaaatagattgaagcaaaataattttaagacaaaaaaaatg aatcaacaatactgagatgatggaaaatgttttaacagatctgcaaagtccgaggaatcgtaataatgttccgatgaagttg ccataccacgtaattctttcatattcaaacctgctgatttgtctgtactgattccaaccgtgtacacaacaacgccttggcttttaa gtgcttcttcgcctgaaatttttcagattctagaaaattttagggtaacgctgaaagagcctaaatttggaacaagtgatttgct ggcacgcacattcaggtgaaaacccgatgacgggttgtgtcccataactatgcccttattaataatacattacggcttaaac atcctagtcagtgtacggtactagaaggcatgaa

### Genotyping protocols

#### Genotyping the *inx-1(tm3524) X* allele

*inx1_tm_F: GCCTGTCAGTTGCCAAATCT*

inx1_tm_R1: GCAGTGTCCATGTTCCATTG

inx1_tm_R2: ATGTGTGTCCAGAAGCGATG

Anneal at 55°C, elongate for 1 min at 72°C. Run amplified products on a 2% agarose gel. Homozygous wild-type will produce 542bp & 159bp bands, homozygous inx-1(tm3524) X will produce a single 304bp band, and a heterozygous inx-1(+/tm3524) X will produce three bands at 542bp, 304bp & 159bp.

#### Primer set for *inx-1* exome sequencing

*ex1_inx1_F: CGGCACGGATGAAGTAGTTT*
*ex1_inx1_R: TTGGTTTCATCTCCCGTTTC*
*ex1_inx1* product: 576bp

*ex2_inx1_F: AGACGGGAGACCTACGTTGT*
*ex2_inx1_R: CCGCAACTTAGCCGATCTTA*
*ex2_inx1* product: 982bp

*ex3_inx1_F: AAACACGCAATTTTCTGCAA*
*ex3_inx1_R: CACACACGCACATCCATACA*
*ex3_inx1* product: 994bp

*ex4_inx1_F: CTGCTGTCTGCTCGCTAATG*
*ex4_inx1_R: ACGTGGTCGGGTTAGATGAG*
*ex4_inx1* product: 250bp

Anneal at 55°C, elongate for 1 min 15 seconds at 72°C. Send resulting products for Sanger Sequencing performed by GENEWIZ (Azenta Life Sciences).

