## Supplementary figures and images for "Specific configurations of electrical synapses filter sensory information to drive choices in behavior"

### Supplementary Figure 1

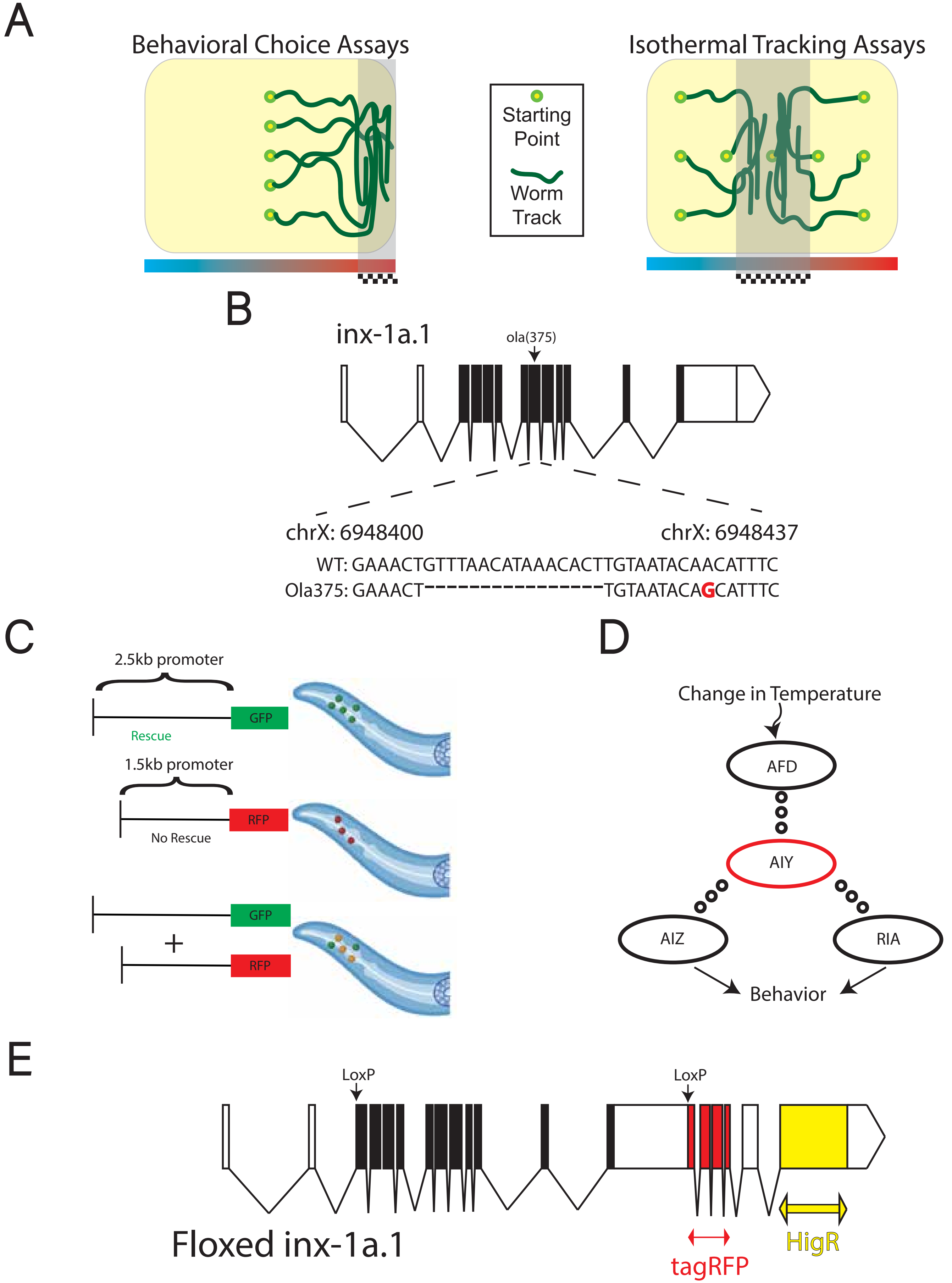

### Supplementary Figure 2

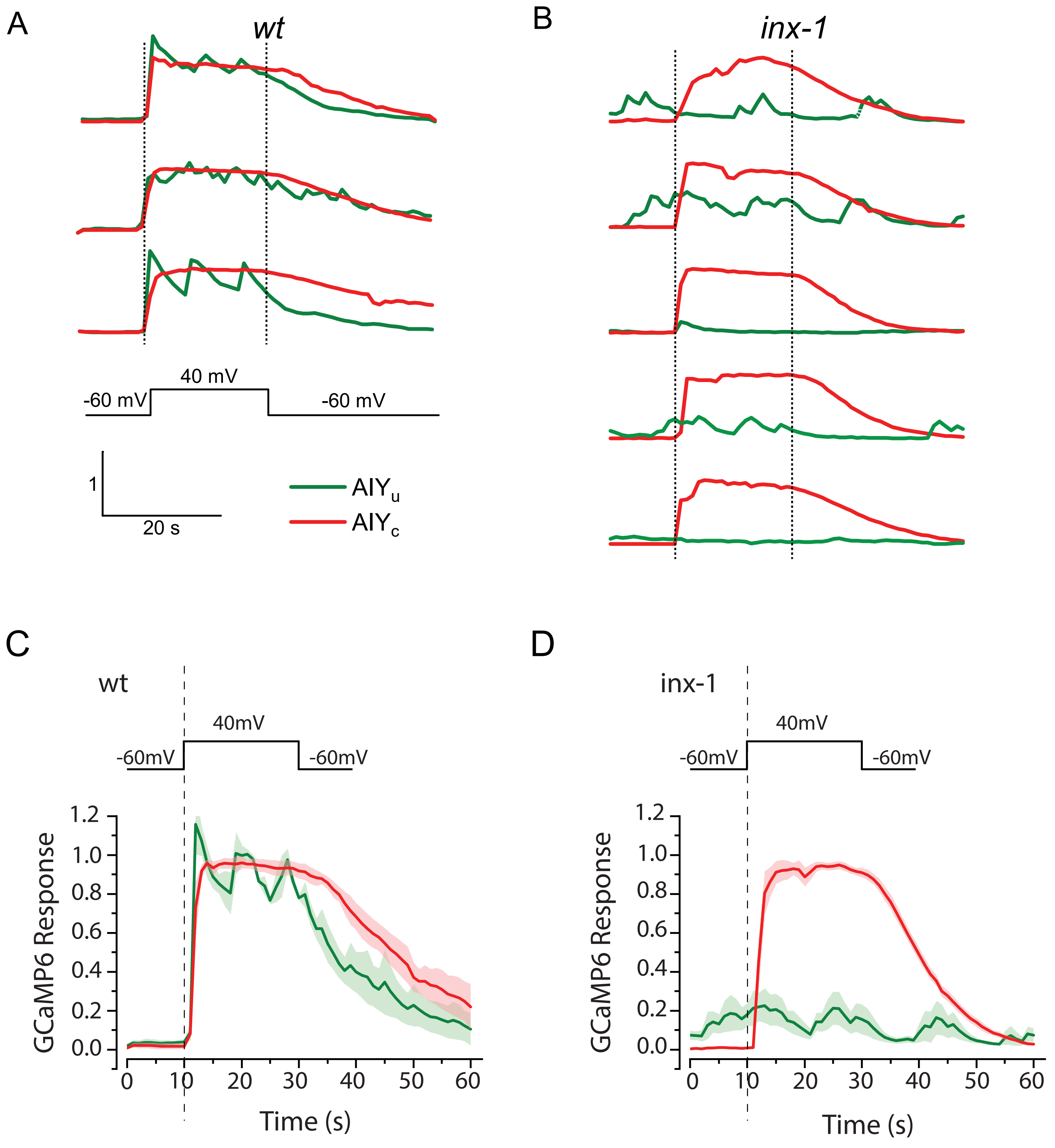

### Supplementary Figure 3

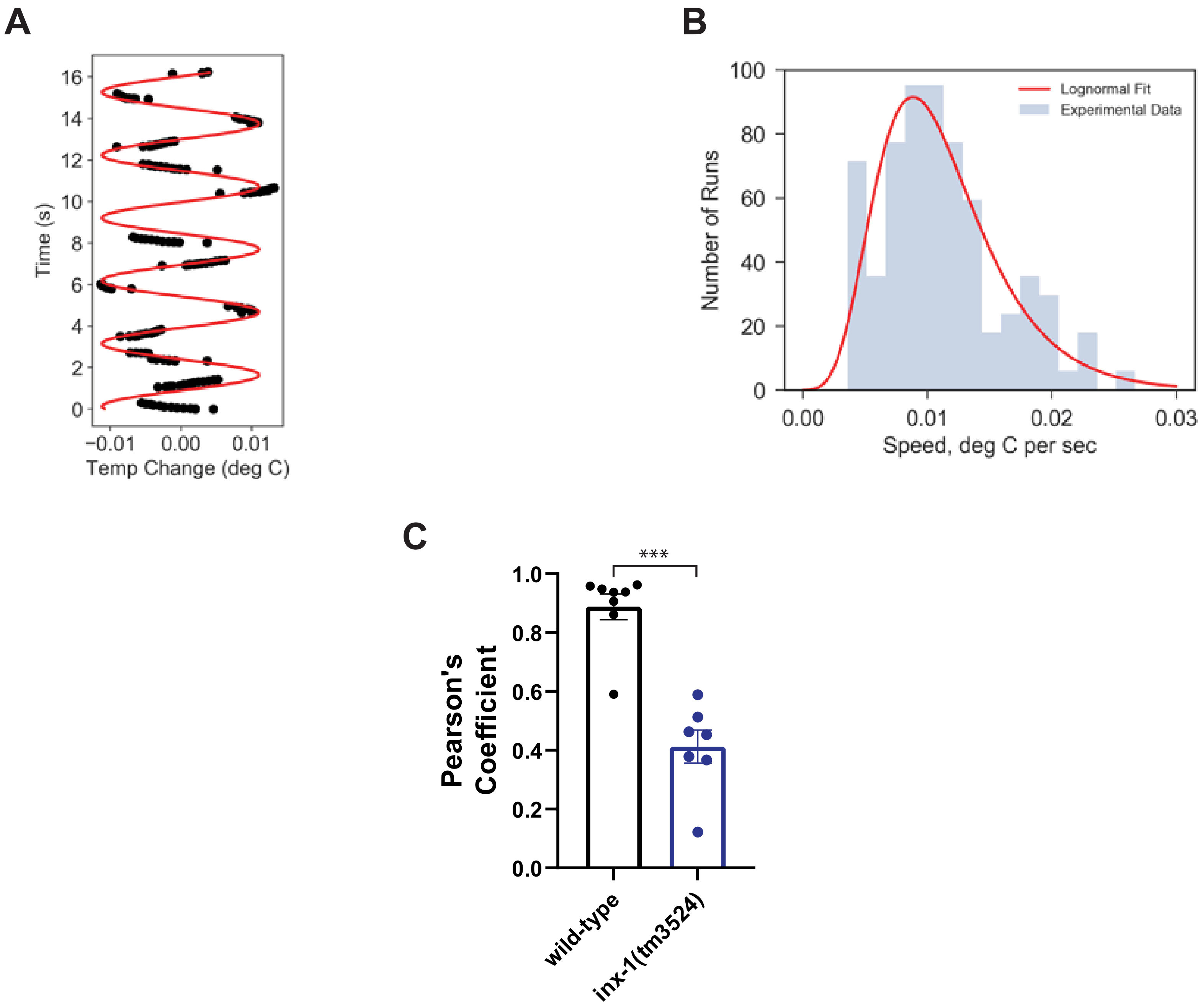
